# Speech auditory-motor adaptation lacks an explicit component: reduced adaptation in adults who stutter reflects limitations in implicit sensorimotor learning

**DOI:** 10.1101/2020.09.06.284638

**Authors:** Kwang S. Kim, Ludo Max

**Author notes:** Now at University of California San Francisco, San Francisco, CA. Correspondence concerning this work should be addressed to Ludo Max, Ph.D., Department of Speech and Hearing Sciences, University of Washington, 1417 NE 42nd St., Seattle, WA 98105-6246.

## Abstract

The neural mechanisms underlying stuttering remain poorly understood. A large body of work has focused on sensorimotor integration difficulties in individuals who stutter, including recently the capacity for sensorimotor *learning*. Typically, sensorimotor learning is assessed with adaptation paradigms in which one or more sensory feedback modalities are experimentally perturbed in real-time. Our own previous work on speech with perturbed auditory feedback revealed substantial auditory-motor learning limitations in both children and adults who stutter (AWS). It remains unknown, however, which sub-processes of sensorimotor learning are impaired. Indeed, new insights from research on upper-limb motor control indicate that sensorimotor learning involves at least two distinct components: (a) an explicit component that includes intentional strategy use and presumably is driven by target error, and (b) an implicit component that updates an internal model without awareness of the learner and presumably is driven by sensory prediction error. Here, we attempted to dissociate these components for speech auditory-motor learning in AWS vs. adults who do not stutter (AWNS). Our formant-shift auditory-motor adaptation results replicated previous findings that such sensorimotor learning is limited in AWS. Novel findings are that neither control nor stuttering participants reported any awareness of changing their productions in response to the auditory perturbation, and that neither group showed systematic drift in auditory target judgments made throughout the adaptation task. These results indicate that speech auditory-motor adaptation relies exclusively on implicit learning processes. Thus, limited adaptation in AWS reflects poor implicit sensorimotor learning.

## Introduction

The neural mechanisms underlying stuttering remain largely unclear, but it has been hypothesized that this disorder of speech fluency may be due to unstable, noisy, or incorrect neural representations (i.e., internal models) of the multiple motor-to-sensory transformations involved in generating acoustic speech output (Cai et al., 2012; Daliri et al., 2013; Hickok et al., 2011; Max, 2004; Max et al., 2004; Neilson & Neilson, 1987; Neilson & Neilson, 1991). Our own research group has postulated that difficulties with the appropriate updating of such internal models would manifest as limited sensorimotor learning when exposed to experimentally altered sensory feedback (Max, 2004; Max et al., 2004). Since that time, a small number of studies have examined sensorimotor learning in individuals who stutter by means of adaptation paradigms in which real-time spectral changes in participants’ auditory feedback lead to compensatory adjustments in speech articulation. Despite using different spectral manipulations (formant shifts that did or did not change the phonological identity of produced vowels), these studies consistently found that the extent of auditory-motor adaptation is limited in adults who stutter (AWS) as compared with adults who do not stutter (AWNS) (Daliri et al., 2018; Daliri & Max, 2018; Kim, Daliri et al., 2020; Sengupta et al., 2016). Although Daliri et al. (2018) did not find adaptation problems in *children* who stutter (CWS) when using an auditory perturbation that changed the produced vowels’ phonological identity (i.e., hearing “bad” when saying “bed”), our own recent study with a perturbation that did not cause such phonological changes yielded an even greater between-group difference in children than in adults. In fact, CWS in the groups that were 3-4 or 5-6 years of age (the groups closest to the onset of their stuttering) exhibited a complete lack of auditory-motor adaptation. Hence, stuttering individuals’ speech auditory-motor learning difficulties are already present at a very young age, and, therefore, have the potential to be fundamentally related to the onset of the disorder (Kim et al., 2020; Max, 2004).

The accumulating evidence for auditory-motor learning difficulties in both CWS and AWS raises the question specifically which sub-processes are affected in stuttering. In studies of upper-limb visuomotor learning, for example, it has been demonstrated that at least two different sub-processes or components contribute to the learning. One is an implicit component (i.e., occurring without learner awareness) that is typically considered to involve the updating of an internal model (i.e., a neural representation of the motor-to-sensory mapping for a given effector system in a given environment). This internal model updating is believed to be driven by sensory prediction errors; that is, a mismatch between the actual sensory consequences and those predicted based on the generated motor command. The second component is explicit (i.e., with awareness of the learner) and involves intentional strategy use. This explicit component is believed to be driven mostly by target error – a discrepancy between movement target and achieved performance. Some of the strongest support for the independence of explicit and implicit components in sensorimotor learning, and for the role of sensory prediction in the implicit component, is found in work by Mazzoni and Krakauer (2006). Those investigators asked participants to make reaching movements while the location of a cursor representing hand position was rotated 45° counter-clockwise around the center of the workspace. When participants were informed about the visual perturbation and instructed to use an explicit strategy to compensate (i.e., aiming 45° clockwise to the target), target error was immediately minimized. Over subsequent trials, however, participants started to reach even further clockwise to the target – thus, target error gradually *increased*. In other words, when target error was already minimized by explicitly adopting an aiming strategy that countered the visuomotor rotation, sensorimotor learning still took place in an implicit manner so as to reduce sensory prediction error (i.e., the discrepancy between planned movement and observed feedback) rather than target error.

Based on those insights, other investigators studying the adaptation of reaching movements have developed experimental methods to estimate the relative contributions of the explicit and implicit learning components. One approach has been to experimentally restrict movement reaction time so that there is insufficient time for the completion of explicit cognitive processes prior to movement onset (Fernandez-Ruiz et al., 2011; Huberdeau et al., 2015). A different approach has been to ask participants to verbally report their intended aiming direction before each trial by naming a number along the circumference of the circular workspace (McDougle et al., 2015; McDougle et al., 2016; Taylor et al., 2014). The reported aiming direction is interpreted as reflecting the participant’s explicit strategy for that trial. The contribution of implicit learning for the trial is computed as the difference between intended aiming direction and actual movement direction. The time course of both learning components across trials has revealed that explicit learning is dominant immediately after introduction of the perturbation but decreases over time whereas implicit learning increases over time (McDougle et al., 2015; McDougle et al., 2016; Taylor et al., 2014).

Converting this limb motor learning paradigm with verbal reporting of the intended movement direction to the domain of speech production is not straightforward. For example, it is not possible to ask naive subjects where in the formant space (i.e., the two-dimensional acoustic space defined by the first two resonance frequencies of the vocal tract) they will “aim” their acoustic output to counteract the perturbation. Naïve speakers know none of the three critical aspects that would be necessary to answer this question: (a) the relative position of different vowels in the acoustic formant space; (b) the intricate relationships between vocal tract postures and acoustic output; and (c) the specific nature of how the experimental perturbation is manipulating their acoustic output in the formant space (see similar comments in Parrell et al., 2017). In fact, due to these complexities, it cannot be ruled out that speech auditory-motor learning is an entirely implicit process. This would, in fact, be consistent with observations based on typical, nonstuttering speakers showing no difference in the amount of adaptation to formant-shifted feedback when instructed to compensate, to ignore the feedback, or to avoid compensating (Munhall et al., 2009). Similarly, in a pitch-shift study with trained singers, the amount of adaptation was not affected by instructions to either compensate or ignore the feedback (Keough et al., 2013). Based on anecdotal information from post-experiment interviews, it has also been reported that none of the participants were aware of making any changes in their speech in response to a formant-shift perturbation (Floegel et al., 2020; Houde & Jordan, 2002). Recently, Lametti et al. (2020) tested task interference when participants simultaneously performed a reach visuomotor adaptation task (manually moving a joystick while seeing misaligned visual feedback) and a speech auditory-motor adaptation task (speaking while hearing formant-shifted auditory feedback). The finding that specifically the explicit component of reach visuomotor adaptation was diminished by simultaneous speech auditory-motor adaptation whereas the tasks did not interfere in the opposite direction was also been interpreted as suggesting that speech auditory-motor adaptation may lack an explicit component (Lametti et al., 2020).

Clearly, there is a need for speech-appropriate paradigms that can be used to quantify any explicit and implicit aspects of speech auditory-motor learning in general, and in individuals who stutter in particular. Determining whether this clinical population’s auditory-motor learning impairment relates to explicit strategy selection, implicit internal model updating, or both, would provide additional insights into both the sensorimotor mechanisms and neural substrates underlying the disorder. Here, we test AWS and AWNS with a novel paradigm developed to obtain initial information regarding participants’ awareness and intent in terms of adaptive articulatory behavior when speaking with formant-shifted auditory feedback. After each trial, participants were asked to report on a visual analog scale to what extent they intentionally changed their speech, and these ratings were used to quantify the explicit component of learning. As a separate innovation, the paradigm also asked participants at regular intervals to select which acoustic stimulus – chosen from a wide range of stimuli that included each participant’s own unaltered productions and various formant-shifted versions thereof – best represented the test words. Given that the perception of speech sounds can change during auditory-motor adaptation tasks (Lametti et al., 2014; Schuerman, Nagarajan et al., 2017; Shiller et al., 2009; but see Schuerman, Meyer et al., 2017, for a contradictory result), the latter addition to the experimental paradigm sought to examine whether reduced speech auditory-motor learning in AWS may be not a true learning problem but a consequence of greater drift in the perceptual target. For example, if one group of speakers were to experience greater target drift in the direction of the auditory perturbation (e.g., an increase in the target first formant [F1] in an experiment with an upward F1 perturbation), their overall extent of adaptation may be limited because smaller adaptive changes are sufficient to achieve the tolerated perceptual distance between heard feedback and intended target.

## Materials and Methods

### Participants

Thirty participants provided informed consent in accordance with procedures approved by the University of Washington Institutional Review Board. This group consisted of 15 adults who stutter and 15 adults who do not stutter, with pairs of stuttering and nonstuttering participants individually matched for age (± 3 years), sex, and handedness. All participants were (a) between 18 and 50 years old, (b) native speakers of American English, (c) without history of speech, language, and hearing disorders (other than stuttering in the case of people who stutter), (d) naive to the purpose of the study, and (e) not taking any medications with a possible effect on sensorimotor functioning at the time of the study. None of the participants had prior experience participating in a speech experiment involving auditory perturbations. All but two participants had binaural hearing thresholds ≤20 dB HL for the octave frequencies 250–4000 Hz. One stuttering participant had a threshold of 25 dB HL for 1 kHz and 4 kHz in both ears, and another stuttering participant had a threshold of 25 dB HL for 4 kHz in the right ear (note that “passing” an adult hearing screening requires ≤25 dB HL tested at 1, 2, and 4 kHz according to the adult hearing screening guidelines of the American Speech-Language-Hearing Association, n.d.). Unfortunately, one stuttering participant’s hearing thresholds could not be determined due to an equipment problem at the time of testing.

The 15 participants from the stuttering group all identified as stuttering, self-reported that the onset of their stuttering had occurred before the age of 8 years, and the presence of stuttering was confirmed by the first author on the day of the experiment. Given that these participants’ study data were collected at the 2018 National Stuttering Association Annual Conference (see section *Experimental setup*), conversational and reading speech samples for severity analysis by a certified speech-language pathologist were recorded later via video call in an effort to limit participants’ time away from educational and social activities at the conference. Unfortunately, this approach failed for two participants who could not be reached after the conference. In addition, one participant’s speech samples did not reach the minimum overall severity score of 10 that is required for the lowest possible classification of *very mild* on the Stuttering Severity Instrument (SSI-4, Riley, 2009). Therefore, the data from this participant and the matched nonstuttering participant were excluded from further analysis.

Two additional pairs of matched participants had to be excluded because the stuttering individuals exhibited dysfluent speech on a large number of trials in the auditory-motor adaptation task. Given that only fluent utterances are appropriate for analysis in an adaptation study (to avoid contamination of the extracted formant data by stuttering-related speech or nonspeech behaviors), there was an insufficient number of trials available for these participants. Consequently, the final data set included 12 individuals who stutter (age *M* = 27.75 years, *SD* = 7.65, range 18-47 years) and 12 individuals who do not stutter (age *M* = 28.08 years, *SD* = 7.45, range 20-48 years). In each group there were 9 male and 3 female participants. Based on self-report, 11 participants in each group were right-handed and one participant in each group was left-handed. Stuttering severity for the participants with known SSI-4 classification ranged from very mild to very severe (2 very mild, 3 mild, 3 moderate, 1 severe, and 1 very severe).

### Experimental setup

Speech auditory-motor adaptation data for the 12 stuttering participants were collected at the 2018 National Stuttering Association Conference in Chicago, IL. Data for the 12 nonstuttering participants were collected in our laboratory at the University of Washington, but care was taken to keep the recording environment as similar as possible to that encountered when recording the stuttering participants on location (i.e., using the exact same equipment, having only the same experimenter present in the room, and not recording in a soundproofed booth).

Participants wore a headset microphone (AKG C544, Shure) placed ∼2.5 cm from the mouth and connected to a USB audio interface (Babyface Pro, RME) operated by a laptop computer (Latitude E5570, Dell). A custom MATLAB (The MathWorks, Inc.) graphical user interface program was used with the Audapter software (Cai et al., 2008; Cai et al., 2010; Cai et al., 2011; Tourville et al., 2013) to implement real-time formant perturbations in the auditory feedback signal. This perturbed signal was routed from the output of the audio interface to a mixer (MMX-24, Monacor) and then presented to the participant via insert-earphones (ER-3A, Etymotic Research). Audapter was used with *downFact* set to 3, *nDelay* set to 5, and a sampling rate of 48 kHz. As recommended in the Audapter manual, we used different online formant tracking parameters for male and female participants by providing the relevant Audapter function (getAudapterDefaultParams.m) with the input “male” or “female” as appropriate.

Total latency of the feedback loop from microphone input to earphones output was 15.3 ms (measured following the procedures recommended in Kim et al., 2020). Prior to each recording session, amplification levels of the audio interface and mixer were calibrated by playing back a speech signal from a loudspeaker and placing the AKG C544 microphone 2.5 cm from the loudspeaker. Amplification levels were then adjusted such that a speech signal measured to be 90.4 dB SPL at that distance of 2.5 cm from the source (i.e., the level that would result in an intensity of 75 dB SPL at a distance of 15 cm from the source, which is our standard calibration procedure for non-headset microphones) resulted in an output intensity of 73 dB SPL in the ER-3A earphones (measured in a 2 cc coupler Type 4946 connected to a sound level meter Type 2250A with Type 4947 1⁄2” pressure field microphone, Bruel & Kjaer). This selected input-output relationship is based on prior work with simultaneous recordings of a speech signal’s intensity at a microphone in front of a speaker’s mouth and at the entrance to that speaker’s ear (Cornelisse et al., 1991).

### Experimental task and data extraction

#### Speech auditory-motor adaptation

A custom MATLAB program asked participants to read monosyllabic words (“bed” and *“*pet”) that appeared on a touchscreen monitor (P2314T or P2418HT, Dell). Words appeared individually, in alternating order. Each word was produced 10 times in a *baseline* phase with unaltered auditory feedback (20 trials total), 60 times in a *perturbation* phase with altered auditory feedback (120 trials total), and 10 times in an *after-effects* phase during which unaltered auditory feedback was restored (20 trials total). In the perturbation phase, the frequency of the first formant (F1) in the participant’s speech was shifted up by 400 cents which caused “bed” and “pet” (vowel /ɛ/) to sound more like “bad” and “pat” (vowel /æ/), respectively (Figure 1). Thus, participants could compensate for the feedback perturbation by pronouncing the words more like “bid” and “pit.” The 400 cents upward F1 shift was applied by setting the Audapter parameter *bRatio* to 1, all 257 elements of *pertAmp* to 1.26 (i.e., a 26% increase), and all 257 elements of *pertPhi* to 0 radian (setting a 0 radian shift in the F1-by-F2 vowel space implements a shift only along the F1 axis).

**Figure 1.**
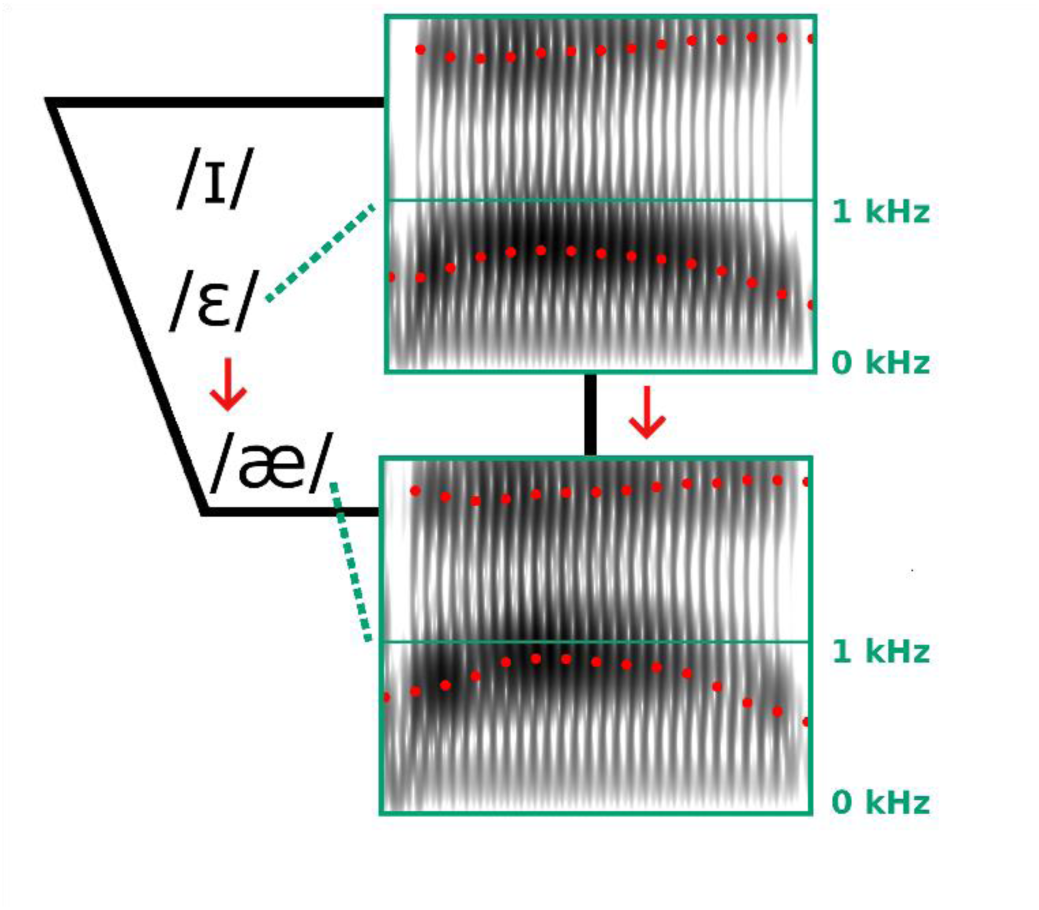
Spectrogram representations of the vowel portion in a participant’s production of “bed” (top) and corresponding heard auditory feedback (bottom) during the perturbation phase of the experiment. Dotted red tracks on the spectrograms indicate the first (F1) and second (F2) formant frequencies. The perturbation shifted F1 up by 400 cents, resulting in the auditory feedback sounding like “bad” which could be compensated by producing “bid.” International Phonetic Alphabet symbols for the vowels in “bid” (/ɪ/), “bed” (/ɛ/), and “bad” (/æ/) are shown in a two-dimensional F1-by-F2 vowel space with F1 increasing from top to bottom.

Participants’ productions were saved directly to computer hard disk for later acoustic analyses. First, to rule out the possibility that one group of speakers may employ a slower rate of speech and thereby benefit from the extra time to make within-trial corrections (rather than truly adapting by adjusting movement planning), we measured each production’s vowel duration with a custom MATLAB program that utilizes Praat algorithms (Boersma & Weenink, 2019). The program automatically marks vowel onset and offset, but we visually inspected each trial and made manual corrections where necessary. We then calculated average vowel duration for each participant. Second, the same software also used Praat algorithms to track the formant frequencies in each production. F1 values for each trial were extracted as the average frequency of this formant across the middle 20% of the vowel (i.e., from 40% to 60% into the vowel). Some trials had to be rejected (8.7% of “bed” and 11.8% of “pet”) for reasons such as mispronouncing the word, yawning, not speaking within the trial’s recording time window, or stuttering. All missing trials were interpolated using four neighboring data points (typically an equal number of preceding and subsequent trials, except for productions that occurred as one of the first or last two trials of the task). For each participant and each word, the extracted F1 values of all trials were normalized by converting the original Hz-based measurements into the relative unit of cents, calculated with the reference value defined as the average F1 frequency across trials 6 to 10 (i.e., the second half of the baseline phase during which no auditory perturbation was applied). Early adaptation was then calculated as the average F1 (in cents) of the first five trials in the perturbation phase. Late adaptation was calculated as the average F1 (in cents) of the last five trials in the perturbation phase.

#### Contribution of explicit strategy use

Throughout the adaptation task, we collected information about the participants’ possible use of an explicit strategy to compensating for the perturbed feedback. After each trial, the MATLAB program obtained information about explicit strategy use by displaying the question “Did you TRY to change your speech? In which direction?” (Figure 2). The participant then moved a slider on the touchscreen monitor to indicate to what extent, if any, they changed their production of “bed” by pronouncing the word “More like bid” or “More like bad” (and similarly for a production of “pet,” whether they pronounced it “More like pit” or “More like pat”). They were instructed to leave the slider in the middle if they had not tried to change their speech. The final slider position was recorded in arbitrary units along the bid-to-bad slider scale (defined as - 100 to +100). For use in the statistical analyses, we calculated for each participant and each test word the average slider position for baseline trials 6 to 10, for the first five trials of the perturbation phase (i.e., the same trials used to define early adaptation), and for the last five trials of the perturbation phase (i.e., the same trials used to define late adaptation).

**Figure 2.**
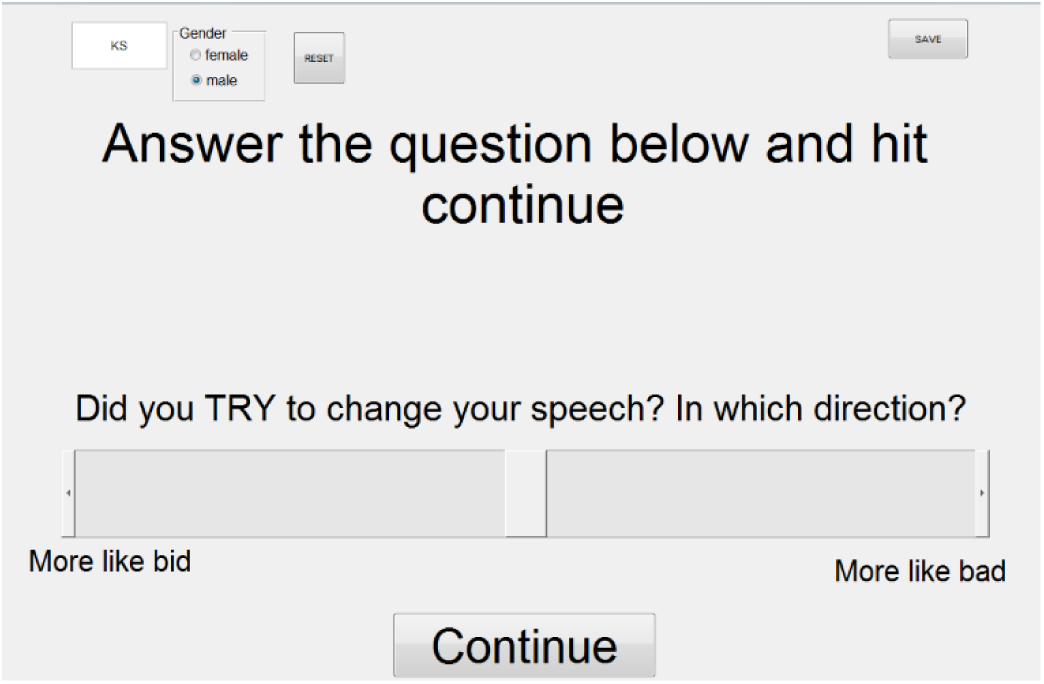
Graphical user interface to collect information regarding participants’ explicit strategy use during the auditory-motor adaptation task. Example shown for “bed” trials. After each trial, the program asked the participant to indicate by means of a slider whether or not they intended to change their speech, and, if so, by how much (i.e., pronouncing the word “More like bid” or “More like bad”). Participants were instructed to leave the slider in the middle of the bar if they did not try to change their speech.

Of course, this post-trial report of intended change is not entirely equivalent to the pre-trial “Where are you going to aim?” question used to estimate explicit strategy use in the limb motor learning literature (e.g., Taylor et al., 2014). However, pilot testing of various potential methods had revealed that it was not possible to ask participants about their intended strategy before the trial (e.g., “How are you going to say bed?”). Even when the pilot test participants were speech-language pathology students with knowledge of formant frequencies and were provided with additional clarification about the intent of the question, they reported being unclear about what was being asked or they believed that it was fundamental frequency rather than formant frequency that they should adjust in order to perceive typical feedback. We therefore examined explicit strategy use only by asking the aforementioned post-trial question about participant intent (“Did you TRY […]”) given that, by definition, only adaptive changes accompanied by awareness constitute explicit learning.

#### Drift in auditory targets

We also collected information regarding potential drift in participants’ auditory target for the test vowel (cf. Shiller et al., 2009). For this purpose, we first pre-recorded each participant producing the words “bed” and “pet” prior to the beginning of the adaptation task. Then at the start of the adaptation task, before the fourth trial of each word, and from then on in steps of ten trials, the MATLAB program displayed the question “What SHOULD bed sound like?” (or “pet”) while also displaying on the same touchscreen monitor a scrollable bar from which auditory stimuli could be selected by double-tapping anywhere along the bar (Figure 3). Double-tapping initiated play-back of the participant’s own pre-recorded production of the test word with or without varying amounts of frequency shift added to F1. In other words, depending on where the participant double-tapped on the scroll bar, the program played back either the original production or different F1-shifted versions of that production. The continuous-scale scroll bar allowed F1 shifts to be selected in the range from −35% to +35% relative to the original production (these extreme ends of the range correspond to −746 cents and +520 cents relative to the original production). Importantly, the auditory stimuli were not simply arranged from lowest to highest F1 along the length of the scroll bar because this would have allowed participants to always select the same auditory stimulus based on visual information and/or memory, even without listing to any of the auditory stimuli (e.g., move to the extreme end on the right side of the scroll bar, then double-tap in the middle of the screen). Instead, we varied with each judgment trial which F1 shift was aligned with the center of the scroll bar. Although this procedure was necessarily associated with a discontinuity in F1 values somewhere along the bar (i.e., when scrolling to the right, F1 values could reach the maximum of +35% and then switch over the minimum of −35% before increasing again), it ensured that participants had to double-tap the screen, listen, judge whether they should scroll to the left or to the right, and decide at which stimulus to stop.

**Figure 3.**
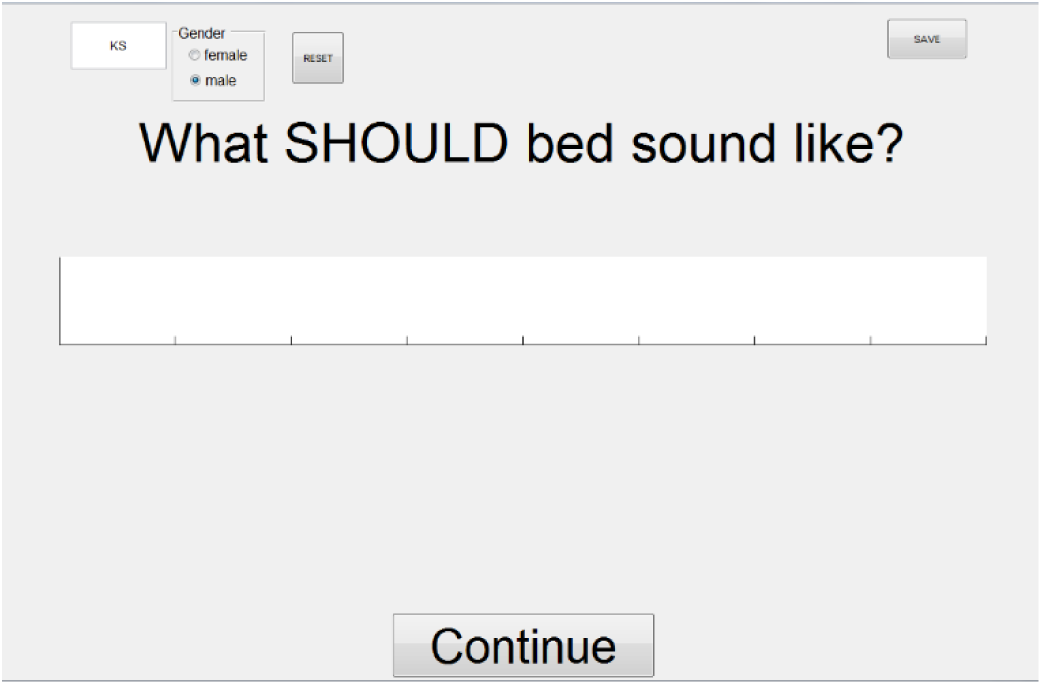
Graphical user interface to collect information regarding participants’ auditory target estimates during the auditory-motor adaptation task. Example shown for “bed” trials. The program asked participants “What SHOULD bed sound like?” and “What SHOULD pet sound like?” The white bar in the center of the screen was scrollable, and scrolling to the left or to the right initiated play-back of a pre-recorded production from the same participant but with increasingly larger downward or upward shifts applied to the first formant. Participants were instructed to always listen to several trials along the scroll bar before making a final decision about which version sounded like the best production of the word.

Prior to the start of the adaptation task, participants were instructed to always listen to multiple stimuli before making their final selection, and to explore the other end of the scroll bar when reaching the end before the best stimulus had been found. Each participant’s final response was recorded as the amount of F1 shift (in cents) applied to the auditory stimulus that was selected as the best representation of the target word. For statistical analysis, we used participants’ selections made in three time windows that corresponded best to those used to define baseline, early adaptation, and late adaptation for the speech acoustics data. Specifically, we averaged the last two selections made before onset of the perturbation to represent baseline (i.e., 2^nd^ and 3^rd^ auditory targets), the first two selections made after onset of the perturbation to represent the early adaptation window (i.e., 4^th^ and 5^th^), and the last two selections made before the end of the perturbation phase to represent the late adaptation window (i.e., 14^th^ and 15^th^).

### Statistical analyses

Using the *ezANOVA* function from the *ez* package (Lawrence, 2016) in R (R Core Team, 2018), repeated measures analysis of variance (ANOVA) was conducted to analyze results for the dependent variables vowel duration and extent of adaptation. For vowel duration, the repeated measures ANOVA tested the Group effect (AWS vs. AWNS), the within-subjects Word effect (“bed” vs. “pet”), and the interaction (Group × Word). For extent of adaptation, the repeated measures ANOVA included the same Group and Word effects but also the within-subjects effect of Phase (early adaptation vs. late adaptation) and all interactions among these variables (Group × Phase, Phase × Word, Group × Word, Group × Phase × Word). For all effects, we report effect sizes as partial omega-squared 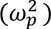 calculated with the *omega.partial.SS.rm* function from the MOTE package in R (Buchanan et al., 2019).

Descriptively, the ratings of intent to change (as a proxy for the explicit component of adaptation) and selection of best target (to examine possible auditory target drift) showed little change throughout the task. We therefore decided to apply independent one-sample *t*-tests to determine if either the AWNS’ or AWS’ ratings and selections deviated from 0 during the baseline, early adaptation, and late adaptation. The *p* values for all six tests that were part of the same family of comparisons (with the intent to change ratings and auditory target selections defined as two separate families), were adjusted with the Holm−Bonferroni method using the *p.adjust* function in R (R Core Team, 2018). Effect sizes for these *t*-tests are reported as Cohen’s *d* calculated with the *cohen.d* function from the *effsize* package in R (Torchiano, 2018).

Lastly, for the group of participants who stutter, we examined whether there was a correlation between extent of auditory-motor adaptation and average stuttering frequency across the conversational and reading speech samples recorded for the SSI-4 (Pearson’s correlation in R; available for only 10 of the 12 AWS for reasons described above).

## Results

### Vowel duration

AWS and AWNS did not differ in vowel duration, F(1, 22) = 1.124, *p* = 0.300, 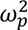 = 0.003. As can be expected based on the consonant context, vowel duration was significantly longer in the word “bed” as compared with “pet,” F(1, 22) = 74.281, *p* < 0.001, 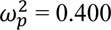, but there was no significant interaction between Group and Word, F(1, 22) = 0.181, *p* = 0.675, 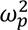 = −0.008.

### Auditory-motor adaptation

Descriptively, both the group of AWNS and the group of AWS showed auditory-motor adaptation, lowering F1 during the perturbation phase relative to the baseline phase (Figure 4). However, the repeated measures ANOVA revealed a statistically significant main effect of Group, *F*(1, 22) = 13.426, *p* = 0.001, 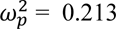, as AWS adapted less than AWNS. The main effect of Phase was also statistically significant (early vs. late adaptation), *F*(1, 22) = 18.417, *p* < 0.001, 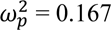, reflecting that late adaptation was more extensive than early adaptation. There was no significant main effect of Word, *F*(1, 22) = 0.149, *p*= 0.703, 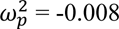. In addition, none of the interactions were statistically significant, Group × Phase, *F*(1, 22) = 1.447, *p* = 0.242, 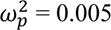, Group × Word, *F*(1, 22) = 0.198, *p* = 0.661, 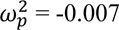, Phase × Word, *F*(1, 22) = 1.565, *p* = 0.224, 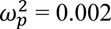, Group × Phase × Word, *F*(1, 22) = 1.105, *p* = 0.305, 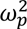 = 0.000.

**Figure 4.**
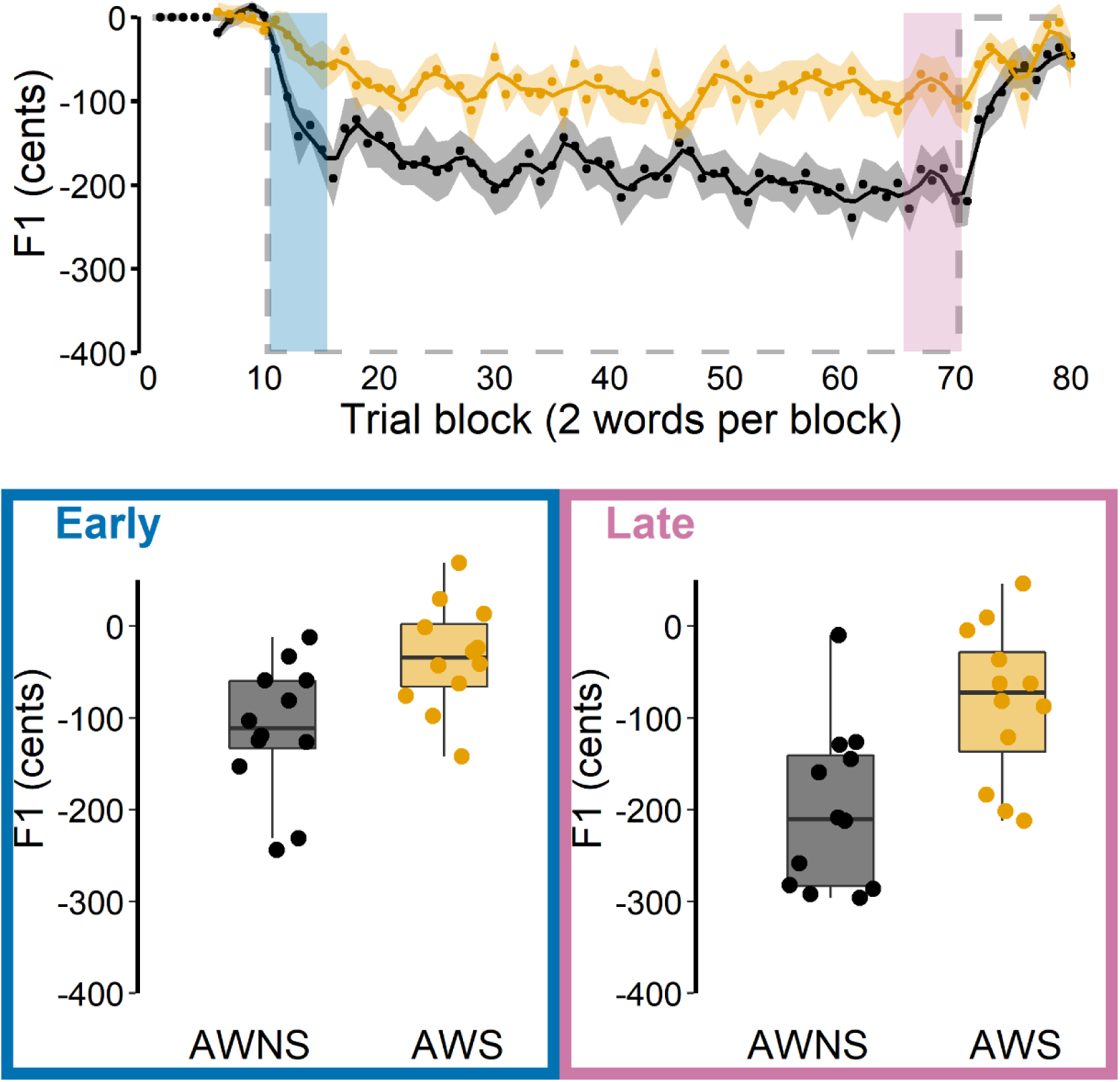
Auditory-motor adaptation for AWS (orange) and AWNS (black). First formant (F1) data are shown in cents, normalized to baseline trial blocks. Top: Group mean F1 for each trial block across the baseline, perturbation, and aftereffect phases; black and orange shaded bands indicate group standard errors of the mean. The grey dashed line represents hypothetical perfect adaptation. The blue shaded area indicates the first five perturbation trial blocks where early adaptation was measured. The pink shaded area indicates the last five perturbation trial blocks where late adaptation was measured. Bottom: Extent of early and late adaptation for each individual participant overlaid on group boxplots.

### Explicit strategy use

Most participants placed the rating slider always at the center of the scale, thereby reporting that they were not trying to change their productions during the task (Figure 5). As expected, during the baseline phase, the ratings did not differ significantly from 0 for either AWNS, *t*(11) = 0.506, *p* = 1.000, *d* = 0.146 or AWS, *t*(11) = −0.641, *p* = 1.000, *d* = 0.185. However, ratings also remained not significantly different from 0 in both the early adaptation phase, AWNS, *t*(11) = 1.399, *p* = 1.000, *d* = 0.404, AWS, *t*(11) = 1.053, *p* = 1.000, *d* = 0.304, and the late adaptation phase, AWNS, *t*(11) = −0.665, *p* = 1.000, *d* = 0.192, AWS, *t*(11) = 0.524, *p* = 1.000, *d* = 0.151. Thus, as a group, neither ANS nor AWS reported any explicit changes in response to the auditory feedback perturbation.

**Figure 5.**
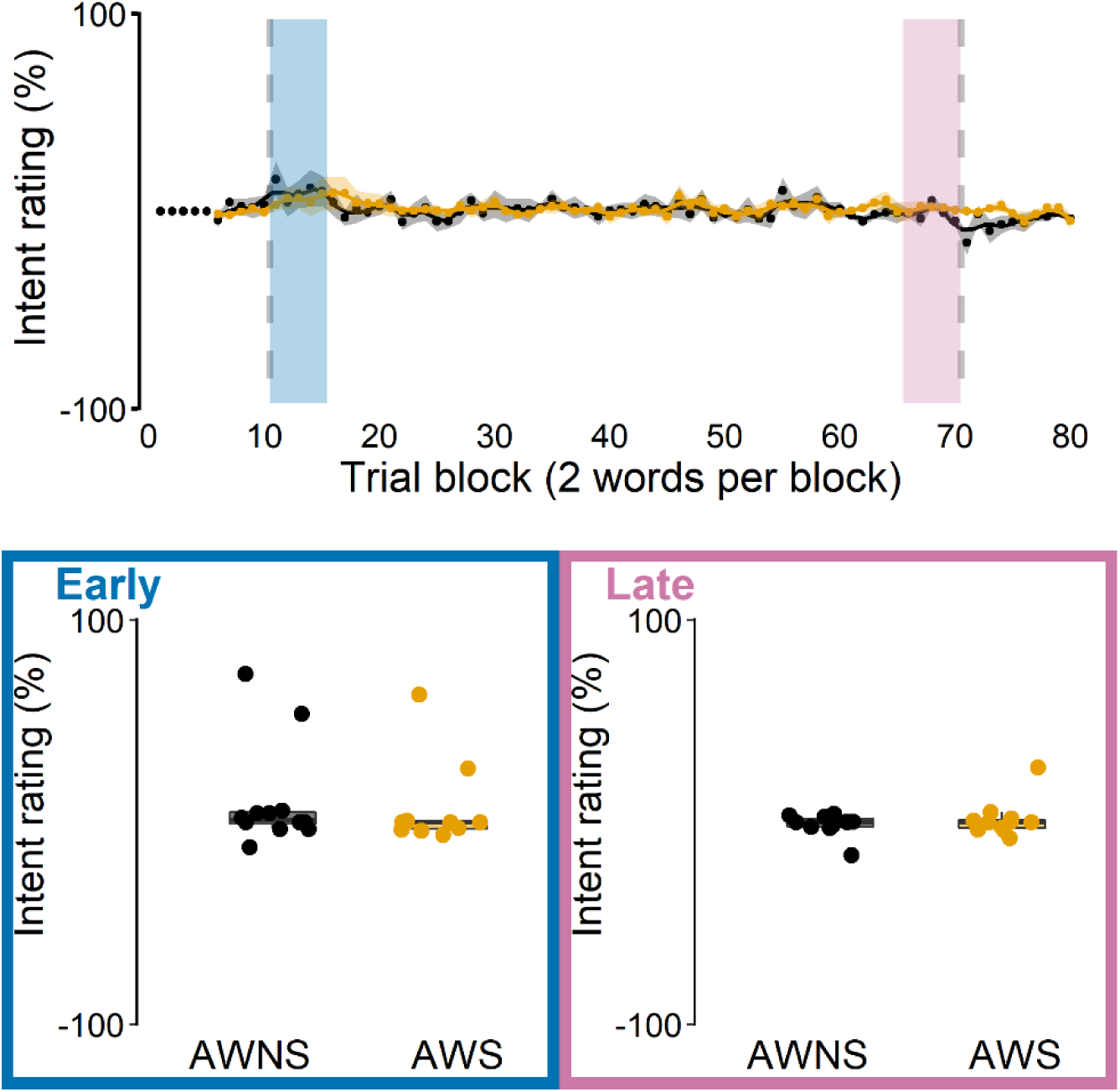
Intent-to-change ratings used to quantify explicit strategy use for AWS (orange) and AWNS (black) during the speech auditory-motor adaptation task. A rating of −100% corresponded to the participant reporting a maximum effort to produce “bed” more like “bid” (or “pet” more like “pit”). A rating of +100% corresponded to the participant reporting a maximum effort to produce “bed” more like “bad” (or “pet” more like “pat”). Top: Group mean intent ratings for each trial block; black and orange shaded areas indicate standard errors of the mean. The vertical dashed lines indicate onset and offset of the perturbation phase. The blue shaded area indicates the first five perturbation trial blocks where intent ratings during early adaptation were measured. The pink shaded area indicates the last five perturbation trial blocks where intent ratings during late adaptation were measured. Bottom: Intent ratings during early and late adaptation for each individual participant overlaid on group boxplots.

It should be acknowledged that a few individuals did move the rating slider away from the center of the scale somewhat consistently. Using an arbitrary threshold criterion that required the mean value of all rating trials in the perturbation phase to deviate more than 10% from baseline in either direction, we found three participants (two from the AWNS group and one from the AWS group) who fit this pattern (Figure 6). However, only one of these participants (AWNS) indicated that they tried to pronounce the test words “More like bid” and “More like pit,” which is consistent with opposing the auditory perturbation and with their speech data indeed showing such adaptive behavior. When asked about these responses after the experiment, this participant actually explained explicitly changing more toward “bid” and “pit” because the auditory feedback sounded too much like “bad” and “pat.” On the other hand, the two other participants (one AWNS and one AWS) both indicated that they tried to pronounce the test words more like “bad” and “pat,” which would exaggerate the auditory error, and which was incongruent with their speech behavior. Hence, it is possible that these participants’ responses were driven by the spectral characteristics of the auditory feedback (given that speech auditory-motor adaptation is always incomplete, feedback does continue to sound more like “bad” and “pat”) rather than by the implementation of an explicit articulatory strategy. Anecdotally, when asked unstructured follow-up questions during debriefing, some participants reported that they did notice changes in vowel *perception* toward “bad” and “pat,” but even those participants denied making any adjustments in their *produced* speech. Thus, with the exception of 1 of 24 participants who was able to precisely describe an explicit strategy involving articulatory changes that opposed the consequences of the auditory perturbation, the perceptual ratings indicate that participants were unaware of the adaptive articulatory adjustments in their speech.

**Figure 6.**
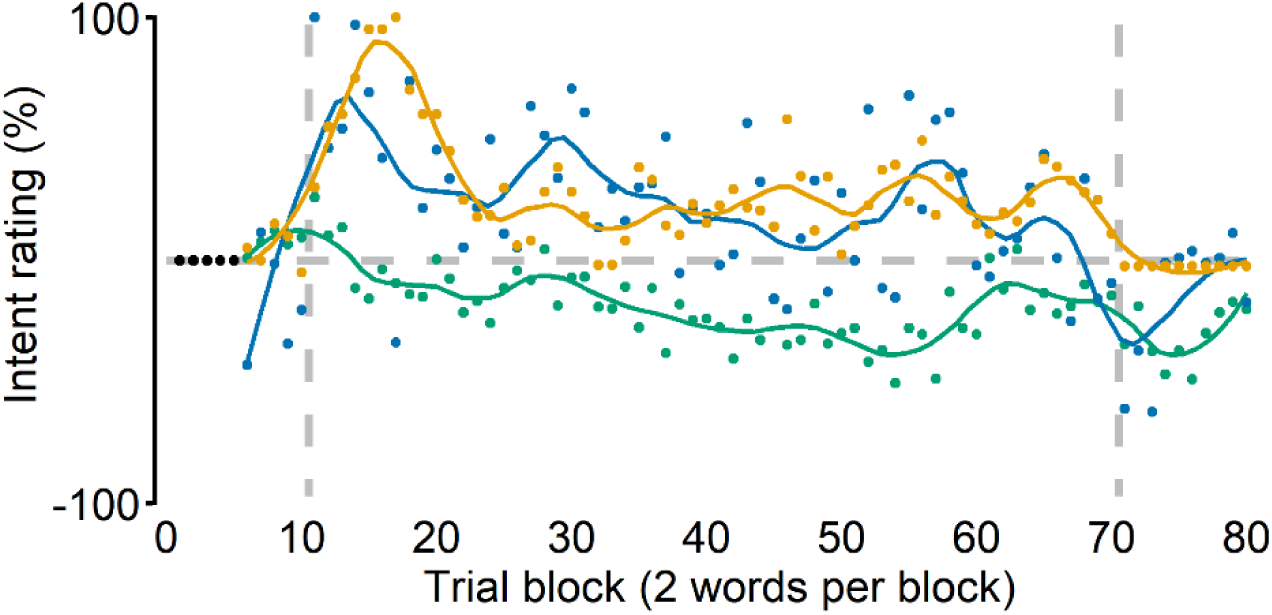
Intent-to-change ratings from the three individual participants (two AWNS and one AWS) who consistently provided non-zero ratings. One AWNS (green) indicated trying to pronounce the test words more like “bid” and “pit,” which is consistent with the direction of change necessary to compensate for the auditory perturbation, and their speech data did show such adaptive behavior. The two other participants (AWNS in blue and AWS in orange) both indicated trying to pronounce the test words more like “bad” and “pat,” which would exaggerate the auditory error, but these ratings were not congruent with their speech behavior. Vertical dashed lines indicate onset and offset of the perturbation.

### Auditory targets

Even in the baseline phase, participants tended to select as the best target (“What SHOULD bed/pet sound like?”) versions of the test words in which F1 had been slightly raised (Figure 7), but these increases in F1 did not significantly differ from 0 cents for either AWNS, *t*(11) = 1.831, *p* = 0.472, *d* = 0.529, or AWS, *t*(11) = 2.355, *p*= 0.229, *d* = 0.680. Most importantly, target selections made during the early adaptation phase after the auditory perturbation was introduced also did not significantly deviate from 0 cents for either AWNS, *t*(11) = 1.110, *p* = 1.000, *d* = 0.320, or AWS, *t*(11) = −0.592, *p* = 1.000, *d* = 0.171. Lastly, this same result of the selected targets’ F1 not being significantly different from 0 cents was also obtained during the late adaptation phase for both AWNS, *t*(11) = 1.040, *p* = 1.000, *d* = 0.300, and AWS, *t*(11) = 0.064, *p* = 1.000, *d* = 0.019. Thus, exposure to the perturbation did not cause either group to select target productions with an increased or decreased F1 relative to the participants’ own pre-test recordings.

**Figure 7.**
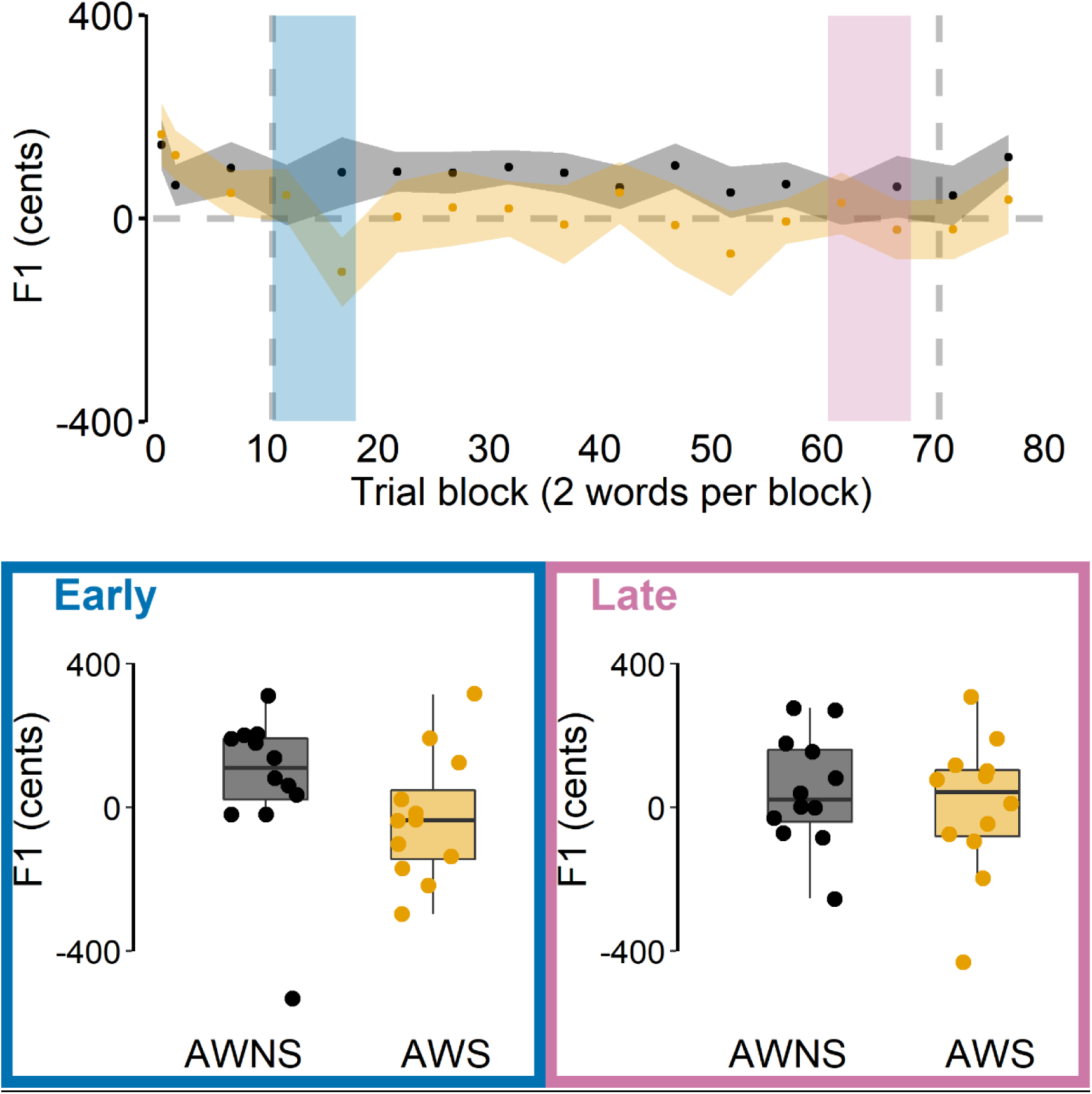
Amount of first formant (F1) shift applied to participants’ own pre-recorded speech in auditory stimuli selected by AWS (orange) and AWNS (black) as the best representation of test words produced throughout the auditory-motor adaptation task. Top: Group mean data and standard errors of the mean. Vertical dashed lines indicate onset and offset of the perturbation. Blue and pink shaded areas correspond to early and late perturbation trial blocks, respectively. Bottom: Stimuli selected during early and late adaptation shown for individual participants overlaid on group boxplots.

### Stuttering frequency and auditory-motor adaptation

For AWS, a stuttering frequency measure averaged across the conversational and reading speech samples from the SSI-4 evaluations was not statistically significantly correlated with either early adaptation, *r*(8) = −0.149, *p* = 0.680, or late adaptation, *r*(8) = −0.246, *p* = 0.493.

## Discussion

In this study, we attempted to dissociate explicit and implicit components of speech auditory-motor learning in stuttering and nonstuttering individuals. Participants produced the monosyllabic words “bed” and “pet” in a baseline phase with unaltered auditory feedback, a perturbation phase during which F1 in the auditory feedback was shifted 400 cents up (causing the words to sound like “bad” and “pat”), and an after-effects phase in which unaltered auditory feedback was restored. To estimate any adaptive changes that were due to explicit strategy use, computer software asked participants after each trial to report by means of a continuous on-screen scale if and how they changed their production of the test word on that trial (i.e., “More like bid/pit” or “More like bad/pat”). To estimate participants’ auditory target for the test word, the software asked every 10 trials “What SHOULD bed/pet sound like?” Participants responded to this question by selecting their preferred version of the word from a range of auditory stimuli that were created by modifying F1 in the participant’s own pre-recorded production of the word from −35% to +35% F1 (including 0% F1 shift).

As a first finding, the individuals who stutter adapted significantly less than the individuals who do not stutter in terms of both early adaptation (i.e., immediately after introduction of the perturbation) and late adaptation (i.e., at the end of the 120-trial perturbation phase). Descriptively, the extent of adaptation observed for the stuttering group was only ∼50% of that observed for the nonstuttering group. This finding again replicates the results of previous studies showing substantial speech auditory-motor learning limitations in adults who stutter (Daliri et al., 2018; Daliri & Max, 2018; Kim et al., 2020; Sengupta et al., 2016). It is worth noting that, in the current study, the number of trials produced during the abrupt-onset perturbation phase (i.e., 120 trials) was greater than the number of perturbation trials in previous studies: Daliri et al. (2018) included 54 trials in their gradual-onset perturbation phase, and 18 of those trials were produced while the perturbation was still being ramped up; Daliri and Max (2018) included 105 trials in their abrupt-onset perturbation phase; Kim et al. (2020) included 90 trials in their abrupt-onset perturbation condition, and although they included 120 trials in their gradual-onset perturbation condition, 60 trials from the latter condition were produced while the perturbation was still being ramped up. Nevertheless, despite this greater number of perturbation trials, stuttering individuals in the present study did not show any benefit of such extended exposure to the auditory feedback perturbation. That is, the difference in adaptation between the two groups did not show any evidence toward decreasing as more trials were performed. Thus, these new results add further support to the growing base of evidence indicating that stuttering is associated with impaired speech auditory-motor learning (Daliri et al., 2018; Daliri & Max, 2018; Kim et al., 2020; Max, 2004; Max et al., 2004). Moreover, the results confirm that prior findings of a between-group difference in this form of sensorimotor learning were not merely due to the use of relatively short perturbation phases that captured only relatively early stages of learning. This strong behavioral evidence of fundamental speech auditory-motor learning limitations in individuals who stutter aligns well with neuroimaging data revealing atypical structural and functional connectivity for auditory-motor integration in the same population (e.g., Kell et al., 2018; Watkins, 2011).

As a second finding, participants’ trial-by-trial reports of whether or not they tried to change their speech indicated an absence of intent to change throughout the entire perturbation phase. Given that both the stuttering group and the nonstuttering group did, in fact, partially adapt to the perturbation (although to a different extent for the two groups), these self-reports of no intent to change confirm that speech auditory-motor adaptation occurs without implementing an explicit strategy to compensate for the perturbation. This finding is fully in agreement with other studies demonstrating that (a) there is no difference in adaptation to pitch-shifted auditory feedback when participants are asked to compensate vs. ignore the feedback (Keough et al., 2013), (b) there is also no difference in adaptation to formant-shifted auditory feedback when participants are asked to compensate, ignore the feedback, or explicitly avoid compensating (Munhall et al., 2009), and (c) the explicit component of reach visuomotor adaptation is diminished by simultaneous speech auditory-motor adaptation but speech auditory-motor adaptation is not affected at all by the simultaneous visuomotor task (Lametti et al., 2020). Combined, our participants’ own direct reports and those prior results based on adaptation tasks with varying instructions provide compelling evidence to conclude that, unlike reach visuomotor adaptation, speech auditory-motor adaptation is entirely implicit for most participants.

A third finding from the present study is that exposure to the F1-shift perturbation and the resulting adaptation did not change participants’ judgment of which version of their own pre-recorded speech (i.e., unaltered or formant-shifted by a varying amount) best represented a typical production of the test word. In other words, the task did not cause drift in participants’ auditory targets for the test words, and this was true for both the stuttering and nonstuttering groups. This observation is interesting in light of the adaptation-induced perceptual effects reported for nonstuttering speakers by Shiller et al. (2009) and Lametti et al. (2014). Shiller et al. (2009) reported that the perceptual boundary between fricative consonants /s/ (as in “see”) and /ʃ/ (as in “she”) shifted during an adaptation task in which feedback for /s/ was perturbed to sound more like /ʃ/. Specifically, this boundary in the region between the intended and perceived sounds was reported to shift toward the perceived /ʃ/ – that is, some stimuli previously identified as /ʃ/ were identified as /s/ after the adaptation task. Lametti et al. (2014) later reported a contradictory result for vowel adaptation: applying an upward F1 perturbation to productions of “head” such that they sounded like “had” caused a perceptual shift not of the boundary between “head” and “had” but of the boundary between “head” and “hid,” with the latter word representing the lower F1 direction in which participants changed their productions as a result of adaptation. Similarly, when F1 in “head” was perturbed downward such that the word sounded like “hid,” there was no change in the boundary between “head” and “hid” but there was a shift of the boundary between “head” and “had.”

In our study reported here, we specifically opted to test for possible drift of the auditory *target* rather than the *boundary* that affects the perceptual classification of ambiguous stimuli in-between two targets. This decision was motivated by four different theoretical and empirical factors: (a) even drift in the specific and narrow classification boundary between two speech sounds may not affect which auditory target a participant is aiming to achieve for each trial; (b) the boundary shifts reported to date were, in fact, not correlated with participants’ extent of adaptation (Lametti et al., 2014; Shiller et al., 2009); (c) there is a long-standing view in speech perception that vowel identification is more continuous and less categorical than consonant identification (Kronrod et al., 2016); and (d) older studies have demonstrated that perceptual boundaries between speech sounds may shift after repeated productions even when feedback is unperturbed or when no feedback is available at all (Cooper, 1979). Clearly, further studies examining both the exact nature of perceptual category boundary shifts and the relationship between perceptual category boundaries and intended perceptual targets will be a worthwhile effort for future work on speech auditory-motor adaptation. Nevertheless, at least in terms of participants’ selection of which auditory stimulus – chosen from a range of stimuli created by F1-shifting each participant’s own pre-recorded speech – represented the best production of each test word, our current study found neither drift during the adaptation task nor a group difference between AWS and AWNS.

Therefore, taken together, the three major findings from this work lead to the conclusion that well-documented limitations in auditory-motor adaptation among individuals who stutter reflect difficulties with *implicit sensorimotor learning*, and that they are not due to differences in either explicit strategy use or perceptual target drift during the adaptation task. One of the remaining challenges for future research will be to determine the implications of this new insight in terms of limitations in specific underlying neural systems and processes. Over the past two decades, the predominant view on sensorimotor adaptation has been that this form of learning reflects the updating of internal forward models that represent the mapping between motor commands and sensory consequences, and it has been theorized that the core problem in stuttering affects the updating of internal models and therefore also the process of making sensory predictions during movement planning (e.g., Hickok et al., 2011; Max, 2004; Max et al., 2004; Neilson & Neilson, 1987). Those theoretical perspectives actually motivated a series of electrophysiological studies that did consistently confirm a lack of motor-to-auditory priming during speech movement planning in AWS as compared with AWNS (Daliri & Max, 2015a; Daliri & Max, 2015b; Daliri & Max, 2018; Max & Daliri, 2019). Thus, there is empirical evidence for a sensory prediction impairment in individuals who stutter, consistent with the previously formulated view that a mismatch between predicted and actual sensory feedback during speech production may cause the central nervous system to apply unnecessary repetitive corrections and interruptions in speech (Max, 2004; Max et al., 2004). What is unclear, however, is whether auditory-motor adaptation tasks such as used in the present study really address mechanisms related to updating internal forward models and generating sensory predictions. Indeed, the debate regarding the fundamental mechanisms involved in sensorimotor adaptation is by no means settled as it has been suggested very recently that implicit adaptation may be driven by direct control policy learning based on prior task errors rather than by the updating of an internal forward model (Hadjiosif et al., 2020). Another recent development stems from the suggestion that the common distinction between explicit and implicit sensorimotor learning itself may be overly simplistic and driven by the use of specific experimental methods whereas the actually relevant processes may not easily map onto those broad categories (Hadjiosif & Krakauer, 2020; Maresch et al., 2020). There is no doubt that a continued strong research focus on sensorimotor learning will be necessary to achieve a deeper understanding of how limitations in the various underlying processes may contribute to the onset of stuttering during childhood speech development.

The findings from this study also have more general implications for some poorly understood aspects of speech auditory-motor adaptation, outside the specific context of stuttering. First, it has been argued in the past that the much more limited extent of auditory-motor adaptation for speech (often in the range 20-40% of the perturbation) as compared with visuomotor adaptation for reaching movements (often in the range 80-95% of the perturbation) might be explained in terms of perceptual adaptation partially offsetting the need for production adaptation (e.g., Houde & Jordan, 2002), discrepancies or trade-offs between the experimentally induced auditory errors and somatosensory errors that arise from adapting one’s typical movements (e.g., Katseff et al., 2012), or physical constraints on the vocal tract’s ability to assume the postures necessary for complete adaptation (Klein et al., 2019). The present study’s confirmation that speech auditory-motor learning constitutes an entirely implicit form of sensorimotor learning suggests, as an additional possibility, that speech adaptation may be less extensive because it lacks the explicit strategy component that contributes substantially to visuomotor learning. For example, Taylor et al. (2014) reported that such an explicit component accounts for approximately one-third of the overall extent of reach visuomotor adaptation. Hence, from this perspective, it would be not surprising to find that an entirely implicit sensorimotor learning task such as producing speech with altered auditory feedback would result in a relatively small extent of adaptation.

A second general implication of the obtained findings relates to the question why speech auditory-motor adaptation is more susceptible to time delays in the auditory feedback signal as compared with reach visuomotor adaptation in the presence of visual feedback delays. It has been demonstrated that even short feedback delays of 100 ms or less greatly reduce the extent of speech auditory-motor adaptation (Max & Maffett, 2015; Mitsuya et al., 2017; Shiller et al., 2020), and that this negative effect cannot be mitigated by prior habituation <u>to the delayed feedback</u> (Shiller et al., 2020). On the other hand, visuomotor adaptation of reaching movements shows only a relatively minor decrease in overall extent of learning even with visual feedback delays of 100-5000 ms (Honda et al., 2012a; Kitazawa et al., 1995; Tanaka et al., 2011), and in at least some conditions this negative effect can be reduced by letting participants habituate to the delay (Honda et al., 2012a; Honda et al., 2012b). In the context of the present work, this differential effect of feedback delay on speech and reach adaptation is interesting because it has been shown recently that the impact on visuomotor tasks is specific to the *implicit* component of learning, with no effect on explicit strategy use (Brudner et al., 2016; Schween & Hegele, 2017). Given that all available evidence indicates that speech auditory-motor adaptation lacks an explicit component, this fundamental difference with reach visuomotor adaptation may be the primary reason for the greater susceptibility to feedback delay during speech sensorimotor learning (Shiller et al., 2020).

Lastly, we acknowledge some methodological limitations of the work reported here. As described above, data for all participants in the stuttering group were collected at the 2018 conference of the National Stuttering Association. A critical benefit of this strategy was that it allowed us to avoid including any participants who had already participated in prior auditory-motor adaptation experiments in our laboratory. However, the total time for informed consent, hearing screening, instructions, microphone and earphones setup, and the experimental task itself was approximately one hour. To address the participants time constraints due to the conference’s educational and social activities, we made significant efforts to keep the overall testing time to a minimum. However, this directly contributed to two methodological limitations. First, as mentioned in the Methods section, the speech samples for stuttering severity analysis were collected at a later date via video meetings, and for two participants no recordings could be made. Second, the auditory stimuli used to test for potential drift in participants’ auditory targets during the adaptation task were generated by spectrally manipulating only a single pre-recorded trial per test word for each participant. For any future replications of this work, it would be recommended to pre-record a large number of trials for each participant, and then create the perceptual stimuli by manipulating the most “representative” trial, for example one that is closest to the participant’s median F1 and F2 values for that particular word.

## Acknowledgements

The authors thank the National Stuttering Association (NSA) Research Committee for accommodating and assisting this research. We are also grateful to Nathalie Farage, Elizabeth Rylance, Elise LeBovidge, and Hantao Wang for help with data extraction and Allie Stewart for conducting stuttering severity assessments. This work was supported in part by grants from the National Institute on Deafness and Other Communication Disorders (R01DC014510 and R01DC17444) and the Canadian Institutes of Health Research (MOP-137001). The content is solely the responsibility of the authors and does not necessarily represent the official views of the National Institute on Deafness and Other Communication Disorders, the National Institutes of Health, or the Canadian Institutes of Health Research.

## Conflict of Interest Statement

The authors declare that there are no conflicts of interest.

## Author Contributions

K.S.K. and L.M designed the experiment and the data analysis pipeline. K.S.K carried out the experiment and analyzed the data. K.S.K. and L.M. wrote the manuscript.

## Data Accessibility Statement

The data that support the findings of this study are openly available in Open Science Framework (OSF) at https://osf.io/s84df/.

## Abbreviations

AWS: Adults Who Stutter
AWNS: Adults Who Do Not Stutter
CWS: Children Who Stutter
F1: First Formant

## Notes

### Competing Interest Statement

The authors have declared no competing interest.

### Summary of Updates

Minor revisions.

